# Functional divergence and potential mechanisms of the duplicate *recA* genes in *Myxococcus xanthus*

**DOI:** 10.1101/766055

**Authors:** Duo-hong Sheng, Yi-xue Wang, Miao Qiu, Jin-yi Zhao, Xin-jing Yue, Yue-zhong Li

## Abstract

RecA is a ubiquitous multifunctional protein for bacterial homologous recombination and SOS response activation. *Myxococcus xanthus* DK1622 possesses two *recA* genes, and their functions and mechanisms are almost unclear. Here, we showed that the transcription of *recA1* (*MXAN_1441*) was less than one-tenth of *recA2* (*MXAN_1388*). Expressions of the two *recA* genes were both induced by ultraviolet (UV) irradiation, but in different periods. Deletion of *recA1* did not affect the growth, but significantly decreased the UV-irradiation survival, the homologous recombination ability, and the induction of the LexA-dependent SOS genes. Comparably, the deletion of *recA2* markedly prolonged the lag phase for cellular growth and antioxidation of hydrogen peroxide, but did not change the UV-irradiation resistance and the SOS-gene inducibility. The two RecA proteins are both DNA-dependent ATPase enzymes. We demonstrated that RecA1, but not RecA2, had *in vitro* DNA recombination capacity and LexA-autolysis promotion activity. Transcriptomic analysis indicated that the duplicate RecA2 has evolved to mainly regulate the gene expressions for cellular transportation and antioxidation. We discuss the potential mechanisms for the functional divergence. This is the first time to clearly determine the divergent functions of duplicated *recA* genes in bacterial cells. The present results highlight that the functional divergence of RecA duplicates facilitates the exertion of multiple RecA functions.

**Author summary:** Myxobacteria has a large-size genome, contains many DNA repeats that are rare in the prokaryotic genome. It encodes bacterial RecA that could promote recombination between homologous DNA sequences. How myxobacteria avoid the undesired recombination between DNA repeats in its genome is an interesting question. *M. xanthus* encodes two RecA proteins, RecA1 (MXAN_1441) and RecA2 (MXAN_1388). Both RecA1 and RecA2 shows more than 60% sequence consistency with *E. coli* RecA (EcRecA) and can partly restore the UV resistance of *E. coli recA* mutant. Here, our results proved their divergent functions of the two RecAs. RecA1 retains the ability to catalyze DNA recombination, but its basal expression level is very low. RecA2 basal expression level is high, but no recombination activity is detected in vitro. This may be a strategy for *M. xanthus* to adapt to more repetitive sequences in its genome and avoid incorrect recombination.

**Highlights:**

1. *M. xanthus* has two *recA*s, which are expressed with significantly different levels. Both *recA*s are inducible by UV irradiation, but in different stages.
2. The absence of *recA1* reduces bacterial UV-irradiation resistance, while the absence of *recA2* delays bacterial growth and antioxidant capacity.
3. RecA1 retains the DNA recombination and SOS induction abilities, while RecA2 has evolved to regulate the expression of genes for cellular transport and antioxidation.

---

RecA, an ATPase recombinase, is the core enzyme for the DNA homologous recombination (HR), as well as a promotion agent for the LexA autolysis in bacteria [1]. The recombination driven by RecA can repair single or double strand DNA (ss or dsDNA) damages, and also the stalled DNA replication fork repaired through the post-replication-repair pathways [2–5]. However, RecA also participates in the chromosome recombination during cell division cycle, in which HR appear between undamaged homologous DNA sequences, resulting in genetic alteration [6–8], and promotes the horizontal gene transfer between different strains [9–12], which also cause genetic diversity. Thus, HR delicately balances the genomic stability and diversity [13–15]. In addition, after binding to ssDNA, the RecA/ssDNA filament complex serves as the signal of DNA damage, resulting in the self-cleavage of LexA, which activates the LexA-dependent SOS response, releasing the LexA-hindered SOS genes. In the best characterized *Escherichia coli* SOS response, LexA autolysis derepresses the expressions of more than 40 genes involving in DNA repair, mutagenesis, and many other cellular processes [1, 16]. Because of its pros and cons in genomic stability and variability, RecA is expressed under strict controls, for example, *E. coli* normally harbors ∼1200 RecA proteins per cell, and increases the RecA expression only after the SOS induction [16].

Most bacteria, such as *E. coli*, have a single *recA* gene, while some bacteria possess duplicate *recA* genes, which, however, have been investigated only in *Bacillus megaterium* and *Myxococcus xanthus* [17, 18]. In the model strain of myxobacteria, *M. xanthus* DK1622, there are two *recA* genes, *recA1* (*MXAN_1441*) and *recA2* (*MXAN_1388*). RecA1 and RecA2 either can partly restore the UV resistance of the *E. coli recA* mutant, and *recA2*, but not *recA1*, was found to be inducible by mitomycin or nidatidine acid [18, 19]. In *B. megaterium*, the duplicate *recA*s were found to be both damage-inducible and similarly showed some DNA repair ability in *E. coli* [17]. It is unclear whether, and how, the duplicate RecA proteins play divergent functions in the DNA recombination and SOS induction.

In this study, we investigated genetically and biochemically the functions of RecA1 and RecA2 in *M. xanthus*. We found that both the *recA* genes were inducible by UV irradiation, but in different periods. While the *recA1* deletion had no significant effect on cellular growth, but reduced the irradiation resistance and the *lexA*-induction ability. Comparably, the absence of *recA2* did not affect the irradiation resistance, but significantly reduced bacterial growth and oxidative resistance. Protein activity analysis *in-vitro* proved that RecA1, not RecA2, had the DNA recombinant activity and was able to promote LexA autolysis. Transcriptomic analysis indicated that the *recA2* gene was crucial for intracellular substance transport and antioxidant activity. We discussed the molecular mechanisms for the functional divergence of RecA1 and RecA2 proteins.

## Results

### 1 Duplicate *recA* genes in *M. xanthus* are both induced by UV radiation

The two RecA proteins of *M. xanthus* DK1622 are highly conserved, and are both homologous to the RecA protein of *E. coli* K12 (EcRecA). The amino acid identity of RecA1 and RecA2 is 64.6%, and either are 59.36% and 62.04% to EcRecA, respectively. Similar to EcRecA [28, 29], RecA1 and RecA2 consist of three structural domains, a small N-terminal domain (NTD), an ATPase core domain (CAD) and a big C-terminal domain (CTD); thereinto CAD contains the conserved ATPase Walker A and Walker B domains and L1 and L2 DNA-binding domains (**Fig. 1A**). The core ATPase domains of RecA1 and RecA2 are highly conserved, while the N- and C-terminal domains are varied. Compared with EcRecA, the two RecAs of *M. xanthus* have more basic amino acids, and the theoretical isoelectric points (pI) of RecA1 and RecA2 are 7.04 and 6.5 respectively; whereas EcRecA is acidic with the theoretical pI of 5.09 (**Fig. 1B**). Differences in amino acid composition suggested that the RecA1 and RecA2 proteins might vary in their functions.

**Figure 1.**
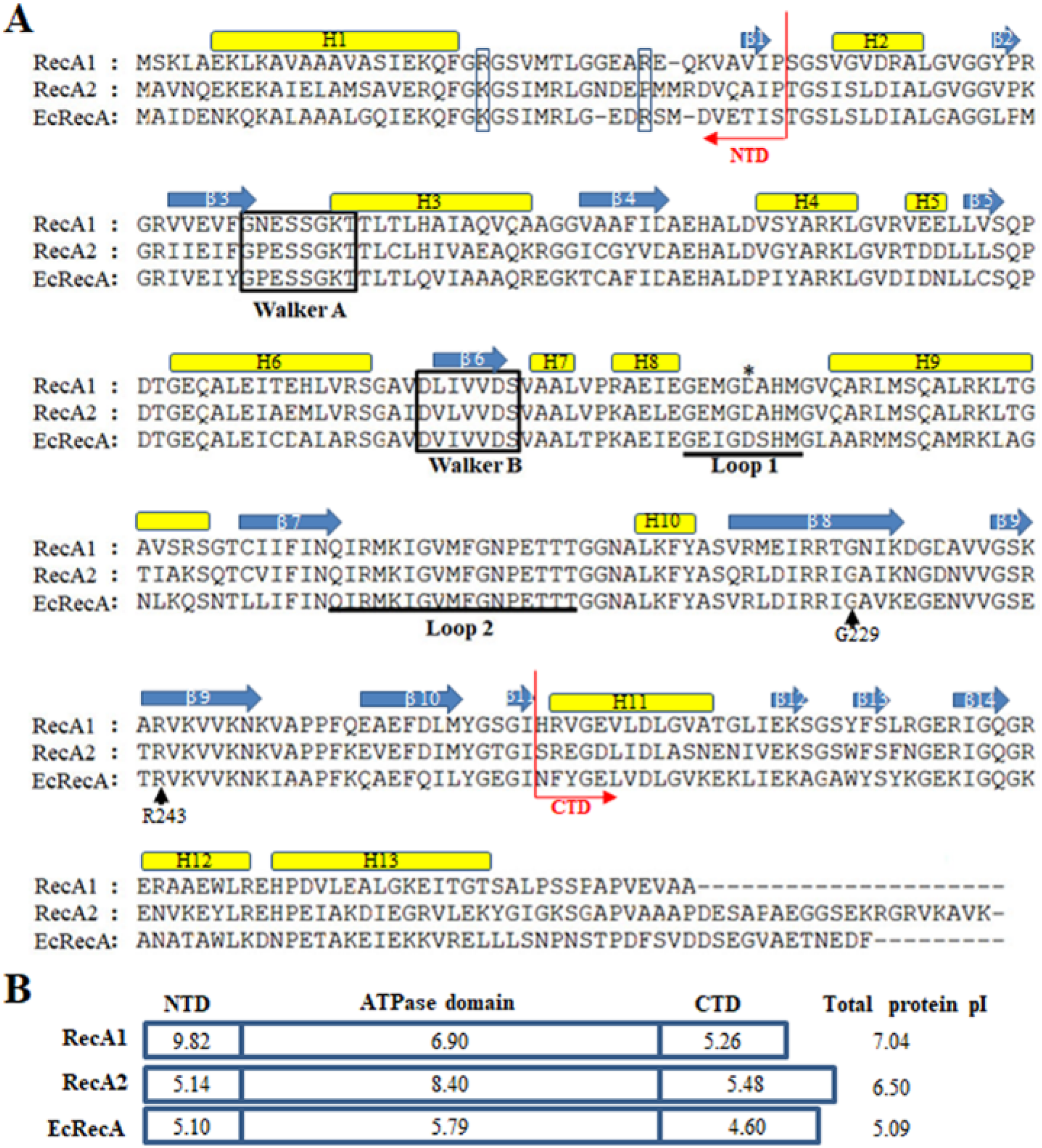
Amino acid sequence comparison of RecAs. (A) Alignment of *M. xanthus* RecA1, RecA2 and *E. coli* RecA (EcRecA, b2699). Positions of the N-terminus (NTD) and the C-terminus (CTD) domains are indicated with red arrows, respectively. Their secondary structures all contain 13 alpha-helixes and 14 beta-sheets, which are indicated above their corresponding amino acid sequences. The ATP binding Walker A and B motifs are marked in frame, and the putative DNA binding sites Loop L1 and L2 are indicated by underlines of the corresponding amino acid sequences. Two reported LexA binding sites (G229 and R243) are indicated by black arrows. K23 and R33 in the N-terminal region of EcRecA are labeled with blue box. (B) The pI features of the domains of three RecA proteins. The theoretical pI values are computed using the ExPASy online tools (Compute pI/Mw).

The SOS response of *M. xanthus* cells on DNA damage can be divided into LexA-dependent and -independent types [19]. The LexA-dependent SOS genes, *e.g. lexA*, typically possess a LexA-box sequence in their promoters. A typical LexA-box sequence was found in the promoter of *recA2*, but not in the *recA1* promoter (**Fig. 2A**). Previous studies reported that *recA2* was obviously induced by naphthylic acid and mitomycin C, but the inducibility was not found in *recA1* by mitomycin C [18, 19]. We treated *M. xanthus* cells with UV irradiation, which directly causes cross-link and single- or double-strand break of DNA, and is a normal induction agent for investigations of bacterial SOS response [30–32]. RT-PCR revealed that *lexA* and *recA2* were up-regulated by 8.3 times and 10.7 times, respectively, after UV irradiation of 15 J/m^2^ dosage (**Fig. 2B**). Interestingly, the *recA1* gene was also UV-induced by 6.4 times. The basal expression level of *recA1* was very low, which was less than one-tenth of *recA2*. The low expression level of *recA1* might be the reason why the expression of *recA1* was not detected by Northern blot [18]. We found that the induction of *recA2* was in the early stage and reached the summit at about 3-hour point after UV treatment, whereas the induction time of *recA1* was delayed and reached the summit until 5 hours after the treatment (**Fig. 2C**). Different expression levels and induction time points implied that the two RecA proteins might involve in different types of DNA damage caused by UV radiation.

**Figure 2.**
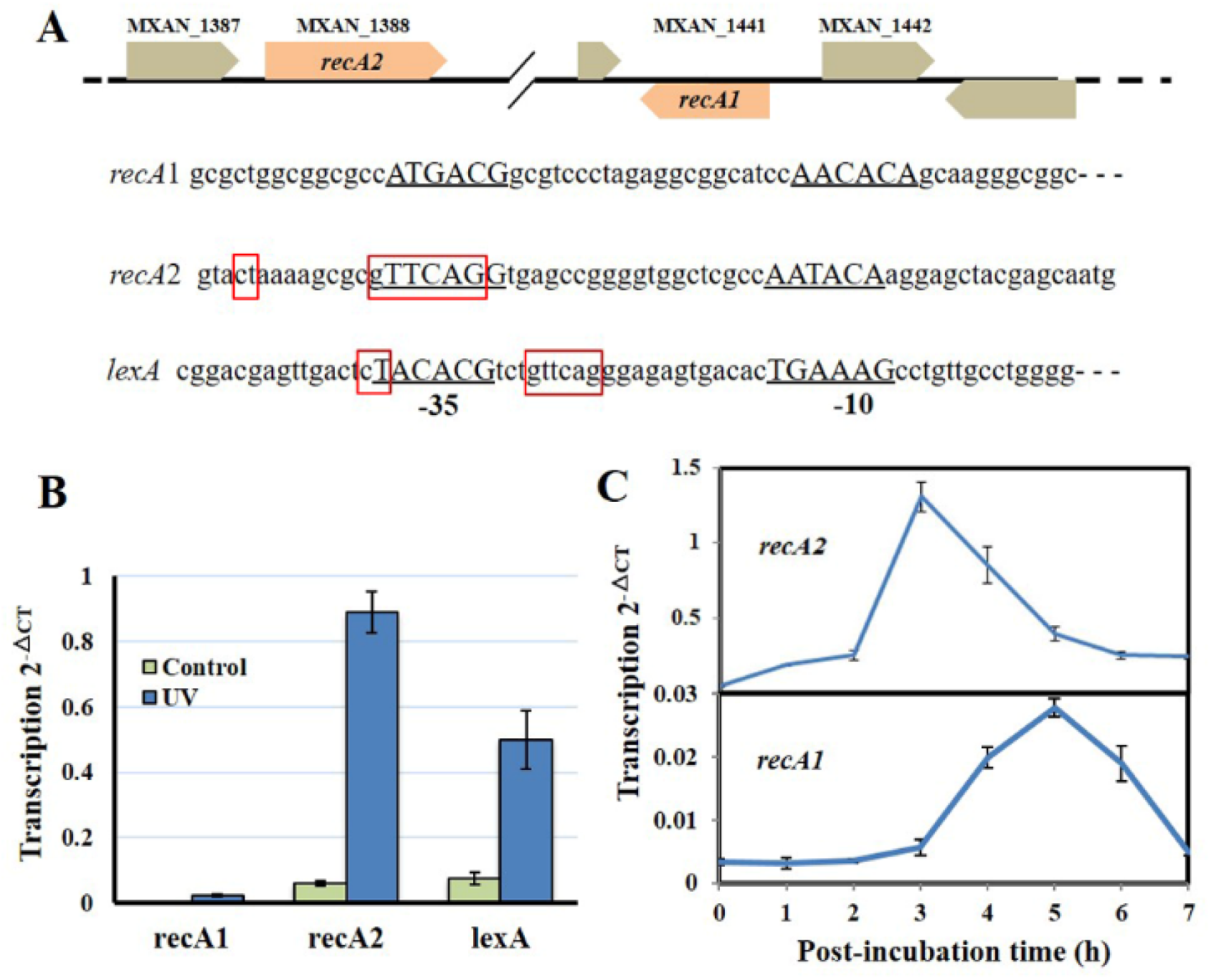
Organization and UV inducibility of the *recA1* and *recA2* genes of *M*. *xanthus* DK1622. (A) Schematic gene location and the promoter alignment of *M*. *xanthus recA1, recA2*. RNA polymerase binding sites (-10 and -35 regions) are underlined and the corresponding nucleotide sequences are in capitals. The SOS box regions are framed in red squares and the sequence in the promoter of *lexA* gene (*MXAN_4446*) was used as a control. (B) UV inducibility of *recA1* and *recA2*, detected with the 4-h cultures after the UV irradiation treatment with the dose of 15 J/m^2^ in a UV crosslinking machine. *lexA* was used as a control. (C) The induction time points of *recA1* and *recA2*. After exposed to UV irradiation at the dose of 15 J/m^2^, the cell cultures were post-incubated at 30 °C, sampled at intervals to extract the total RNA for RT-PCR. The error bars in B and C represent means ± SEM (n = 3, p<0.05 versus inner reference).

### 2 *recA2* plays more important roles than *recA1* in the damage repair process for the growth of *M. xanthus* cells

In previous studies, the *recA2* deletion mutants were not obtained in either *M. xanthus* or *B. megaterium* [17–19]. However, in this study, we successfully obtained the deletion mutant of either *recA1* or *recA2* in *M. xanthus*, named RA1 and RA2, respectively (**Fig. 3A**). According to the employed method, the acquisition probability from reverse screening was ∼10^−6^ for the deletion of *recA1*, but was ∼3.3 × 10^−10^ for the *recA2* deletion. The *recA1* deletion had no significant effect on cellular growth, but the deletion of *recA2* caused the mutant to have a long lag phase, but after the lag phase, growth of the RA2 mutant did not slow down significantly in the logarithmic phase and reached the similar summit as the wild type DK1622 (**Fig. 3B**).

**Figure 3.**
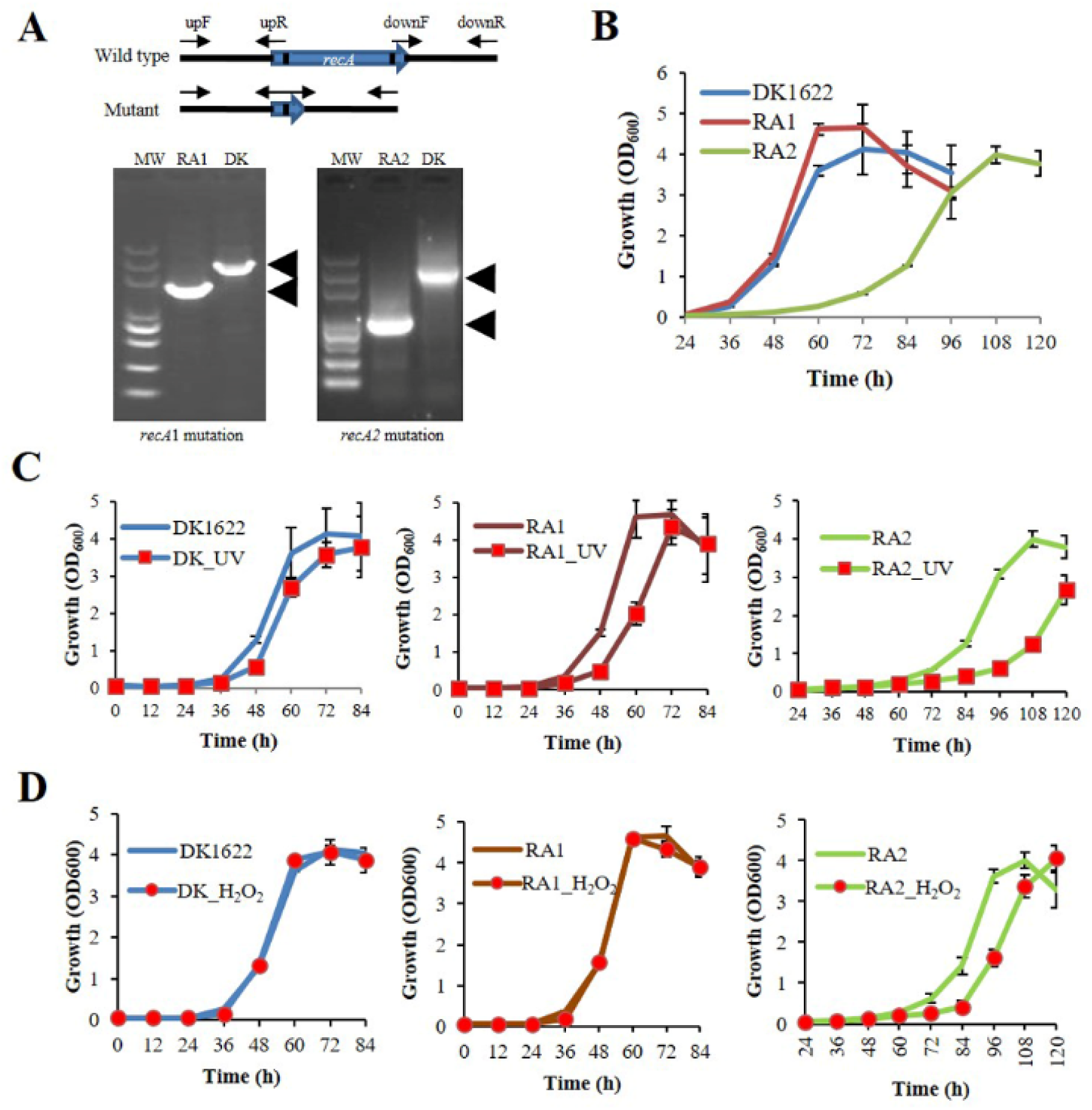
Mutations of *recA1* and *recA2*, and their effects on the growth of *M. xanthus*. (A) Deletion of *recA1* or *recA2* in *M. xanthus* DK1622, using the markerless knockout plasmid pBJ113, producing the RA1 or RA2 mutants. The deletion was verified by PCR using their primer pairs, and sequencing. (B) The growth curves of the *recA* mutants and the wild type strain DK1622 without the treatment of UV irradiation or hydrogen peroxide (H_2_O_2_). (C) Separate growth comparisons of DK1622, RA1 and RA2 with and without the UV treatment at the dose of 15 J/m^2^. (D) Separate growth comparisons of DK1622, RA1 and RA2 with and without the H_2_O_2_ treatment at final concentration of 3 mM for 15 min. The error bars indicate the SEM for six replicates.

UV irradiation majorly causes the DNA damage, while hydrogen peroxide produces oxidative pressure, which damages many kinds of macromolecules, including DNA, leading to the antioxidation response [33]. When treated with 15 J/m^2^ UV irradiation, the growth abilities, compared with that without UV treatment, were all delayed in DK1622, RA1 or RA2, and the growth delay was more serious in RA2 (**Fig. 3C**). When treated with 3 mM H_2_O_2_ for 15 min, DK1622 and RA1 showed almost the same growth curve, while the growth of RA2 was delayed significantly, compared with that of the strains without the treatment (**Fig. 3D**). The results demonstrated that *recA2*, but not *recA1*, is an also crucial factor for the repair of UV irradiation and oxidation damages.

### 3 *recA1* and *recA2* are separately crucial for UV resistance and antioxidation

We measured the survival rates of the wild type strain and the *recA* deletion mutants treated with different dosages of UV radiation (0-25 J/m^2^) and hydrogen peroxide (1-5 mM). All the three strains decreased their survival rates with the increase of UV radiation or H_2_O_2_ dosage. Interestingly, the survival rate of RA1 decreased more significantly than that of RA2 at each UV-irradiation dosage, which, however, had a highly similar survival curve as the wild type strain (**Fig. 4A**). Whereas, the survival rate of RA2 decreased more significantly at each H_2_O_2_ concentration than that of RA1 and DK1622, which showed similar survival curves when treated with hydrogen peroxide (**Fig. 4B**). Thus, RecA1 is probably the key protein for the survival of *M. xanthus* cells under UV irradiation, which is similar to EcRecA [34]; whereas RecA2 involves in the repair of oxidation damage in cell and is important for the survival in the antioxidant process.

**Figure 4.**
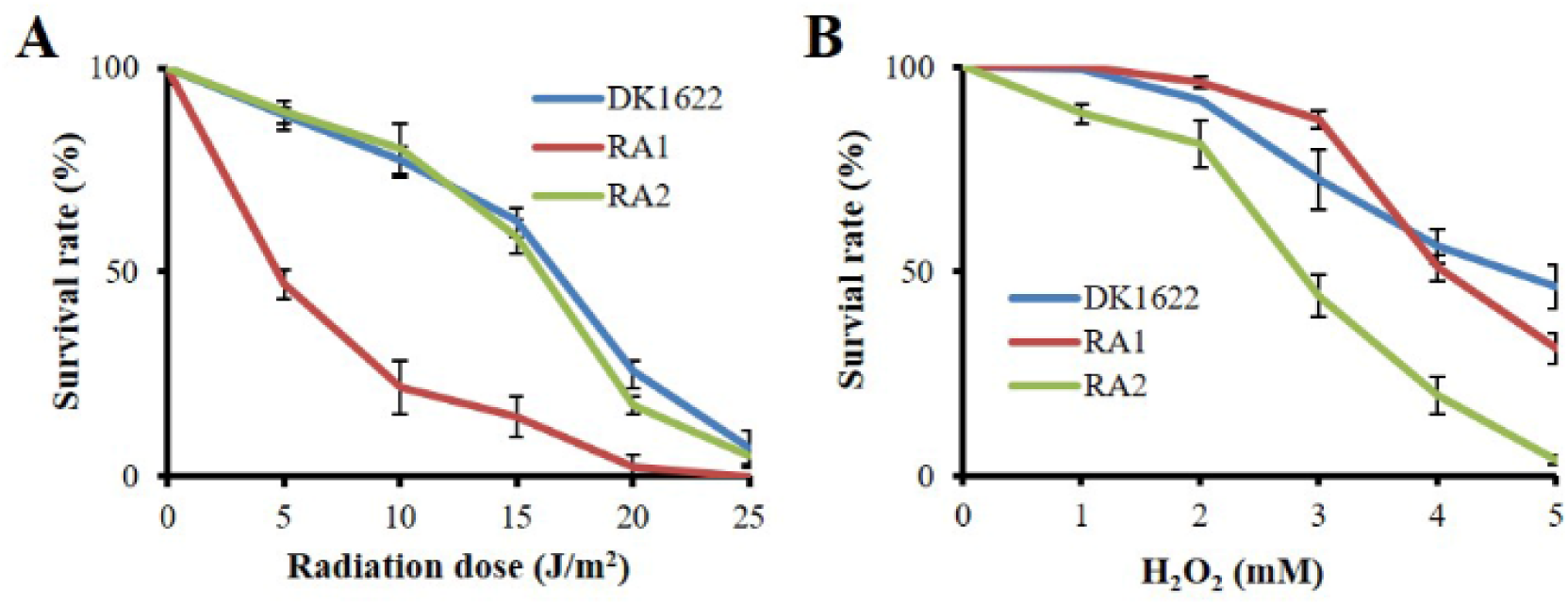
Survivals of *M. xanthus* wild-type strain DK1622 and the mutants RA1 and RA2. (A) Survival curves after the exposure to UV irradiation with different dosages (0-25 J/m^2^). (B) Survival curves after hydrogen peroxide treatment at different concentrations (0-5 mM). The percentage of surviving cells was calculated by comparing with the corresponding non-treated cells. The error bars indicate the SEM for six replicates.

### 4 RecA1, not RecA2, is responsible for HR and LexA-dependent SOS induction

DNA HR and SOS induction are the two main cellular functions of the RecA proteins [1]. We analyzed the *in-vivo* integration abilities of an antibiotic resistance gene into the genomes of DK1622, RA1 and RA2 strains. Calculated from the appearance of resistant colonies, the recombination rate of RA1 was significantly lower than that in either DK1622 (p=0.0088) or RA2 (p=0.0157); while the difference between the recombination rates of RA2 and DK1622 was not significant (p = 0.1049) (**Fig. 5A**). The result determined that *recA1* is crucial for the recombination process in *M. xanthus*.

**Figure 5.**
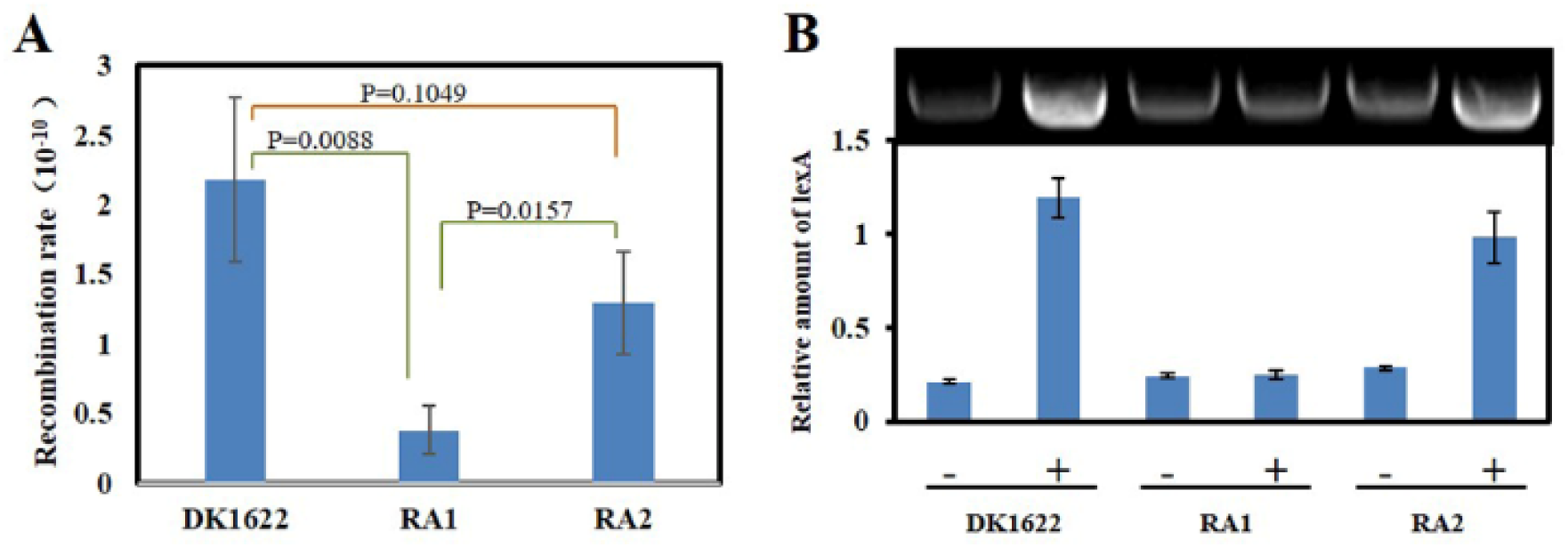
Intracellular DNA recombination rate and induction analysis of *lexA* gene. dependent SOS. (A) The cellular recombination rate of DK1622, RA1 and RA2. The recombination rate was determined by assaying the homologous recombination of an inserted resistance gene in genome. (B) The inducibility of *lexA* gene. Myxococcus *lexA* is a known SOS gene induced through LexA-dependent SOS response and herein, its UV inducibility represents the activation of LexA-dependent SOS reponse. The strains were irradiated with 15 J/m^2^ UV irradiation (+) or mock treatment (-), and the transcription of *lexA* was determined by RT-PCR.

Previous studies indicated that the expression of *lexA* is induced by LexA-dependent SOS response in *M. xanthus* [18]. We compared the transcriptions of *lexA* in *M. xanthus* DK1622, RA1 and RA2 strains in response to the 15 J/m^2^ UV-irradiation treatment. The RT-PCR results showed that *lexA* exhibited the UV inducibility in either DK1622 or RA2, but not in the RA1 mutant (**Fig. 5B**). Thus, the deletion of *recA1*, rather than *recA2*, affected the UV-induction of *lexA*, *i.e.*, RecA1 is responsible for the LexA-dependent SOS induction.

### 5 RecA1 and RecA2 both have the ss- and ds-DNA promoted ATPase activities

We further expressed and purified RecA1 and RecA2 proteins (**Fig. 6A**), and measured their *in*-*vitro* ATPase activities by the quantitative analysis of inorganic phosphorus released from the ATP hydrolysis (**Fig. 6B**). In the reaction mixture without the addition of DNA, RecA1 and RecA2 both exhibited low ATPase activities, and the ATPase activity of RecA2 was some higher than that of RecA1. For example, a microgram of purified RecA2 released 0.1428 nanomole Pi in an hour, which is approximately 2.44 times the hydrolysis capacity of RecA1 on ATP (0.0586 nmol Pi/μg/h). The addition of DNA, especially ssDNA, markedly promoted the ATPase activity of both RecAs, which is consistent with the functionality of the classic RecA proteins [1,13,15]. Thus, RecA1 and RecA2 are both DNA-dependent (more dependent on ssDNA) ATPase enzymes. In the presence of DNA (dsDNA or ssDNA), the ATPase activity increase of RecA1 was higher than that of RecA2. For example, the ATPase activity of 1 ng RecA1 increased by 10.69 times (from 0.0586 to 0.6265 nmol Pi/μg/h) with the addition of ssDNA, while the increase of RecA2 was only twice (from 0.1428 to 0.2857 nmol Pi/μg/h). Similarly, the addition of dsDNA increased the ATPase activities of RecA1 and RecA2 by 6.89 times (from 0.0586 to 0.4038 nmol Pi/μg/h) and 1.86 times (from 0.1428 to 0.2658 nmol Pi/μg/h), respectively.

**Figure 6.**
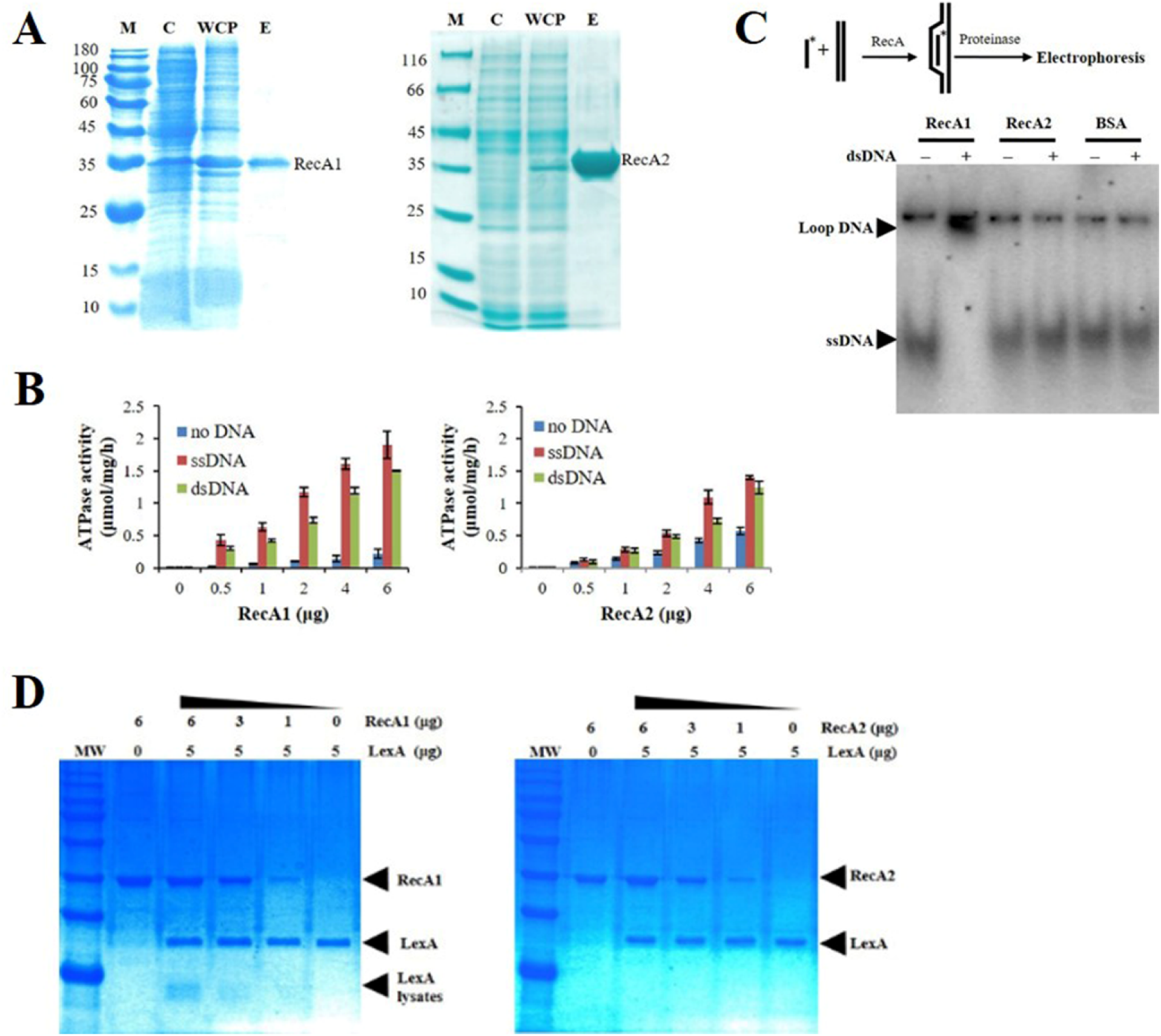
Expression and activity analysis of RecA proteins. (A) Expression and purification of RecA1 and RecA2. M, marker; C, control; WCP, whole cell protein; E, eluent of purified protein. (B) Assays of ATPase activities. The ATPase activity was determined by measuring free phosphate ion (Pi) released from enzymolysis of ATP. The error bar is calculated from three independent repeats. (C) D-loop assay. A 60-nt ^32^P-labeled ssDNA fragment and a superhelical dsDNA (RF M13) sequence were mixed and incubated with and without the addition of purified RecA1 or RecA2 proteins. If the protein has the HR activity, the homologous pairing reaction will be initiated, thus forming the ssDNA-dsDNA complex. Bovine serum albumin (BSA) was used as a control. (D) The promotion ability of RecA1 (left) or RecA2 (right) on the cleavage of LexA proteins. The MxLexA protein was incubated with gradient concentrations of RecA1 or RecA2 proteins in the presence of ssDNA and ATP. Reactions were stopped and visualized on a 1.2 % SDS-PAGE gel stained with coomassie brilliant blue.

### 6 RecA1, not RecA2, has *in-vitro* HR capacity and activates LexA autolysis

Strand assimilation or D-loop formation is a pivotal step in homologous recombination, and is one of the most common biochemical assays for characterizing the activity of RecA-type recombinase [1,13,15,26]. We analyzed the *in*-*vitro* recombination activities of RecA1 and RecA2 in a DNA strand recombination reaction system containing ^32^P-ssDNA and homologous plasmid dsDNA. An obviously-hysteretic radiolabeling band appeared in the lane containing purified RecA1, but not RecA2 (**Fig. 7A**), which determined that RecA1, but not RecA2, has the homologous recombination activity in *M. xanthus*. The result is consistent with the *in-vivo* recombination result (**Fig. 5A**).

**Figure 7.**
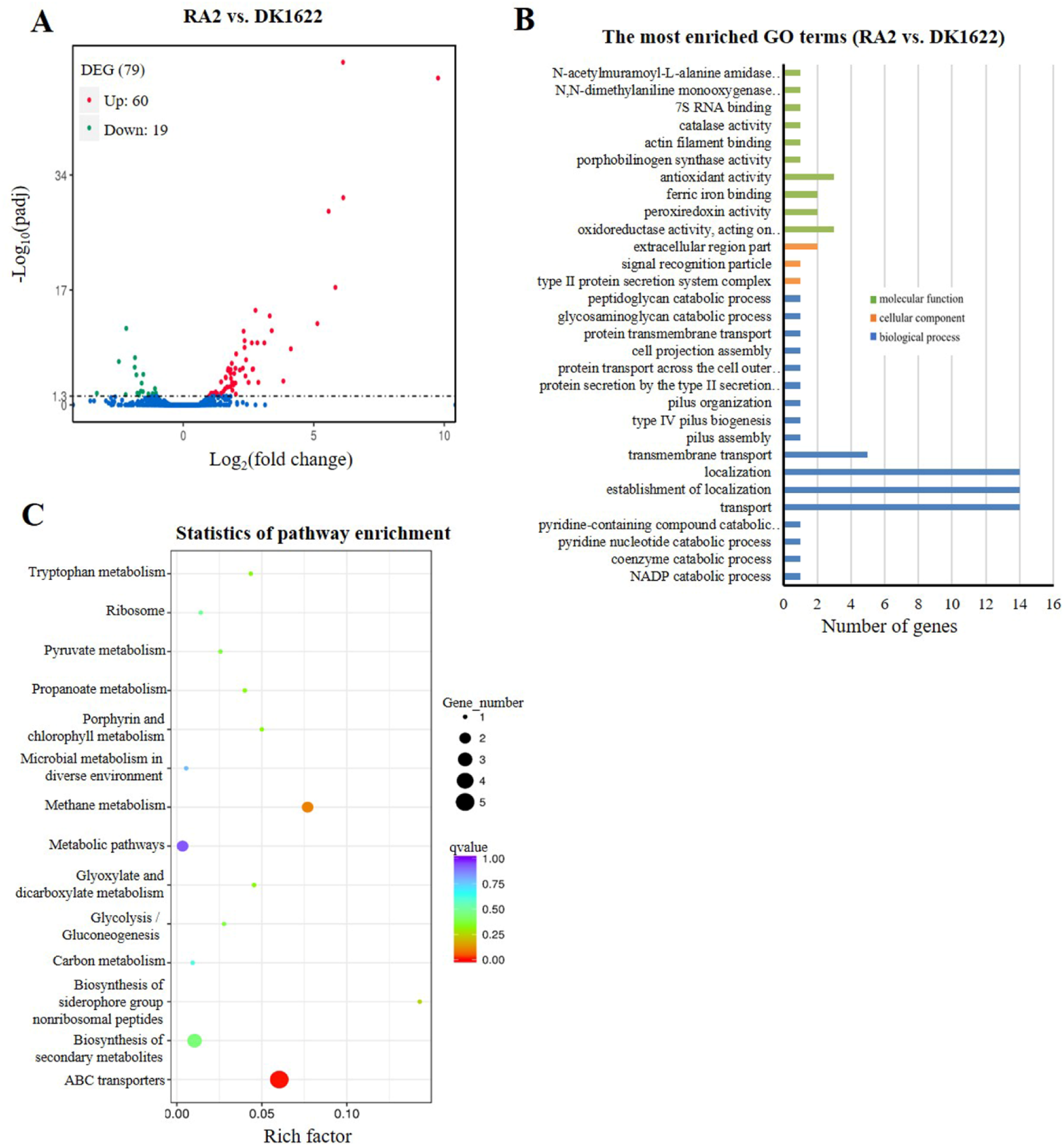
Comparison of transcriptomes between the *recA2* mutant (RA2) and the wild type strain (DK1622). (A) Volcano plot of differentially expressed gene (DEG) distributions. Red dots and green dots represent the up- and down-regulation genes with the significant differences, respectively (padj<0.05). The blue dots represent the genes that have not changed significantly. (B) The distribution of GO category and (C) pathways of DGEs between RA2 and DK1622. The enriched GO are shown in three categories: biological process (blue), molecular function (green), and cellular component (orange).

RecA promotes the LexA autolysis at the A-G peptide bonding sites, thereby enabling the expression of SOS genes inhibited by LexA [35, 36]. We detected the proteinase activity of RecA1 and RecA2 using *M. xanthus* LexA protein as substrate. The results showed that the LexA autolysis fragments were detected in the reaction with RecA1, but not with RecA2 (**Fig. 8**). Thus, RecA1 participated in the LexA-dependent SOS induction reaction, which is also consistent with that RA1 mutant lost the induction ability of SOS gene *lexA* (**Fig. 5B**).

### 7 RecA2 involves in gene regulation for cellular transport and antioxidation

The above genetic and biochemical experiment results demonstrate that RecA1 and RecA2 are both DNA-dependent ATPase enzymes; but RecA1 possesses the functions of classical RecAs, *i.e.* HR capacity and LexA-cleavage promotion ability, while its duplicate RecA2 is divergently evolved to the function in damage repair for growth and antioxidation. To investigate the potential mechanisms of RecA2 in *M. xanthus*, we compared the transcriptomes of the *recA2* mutant (RA2) and the wild type strain DK1622. Totally, 79 genes were found to be expressed differentially (Padj<0.05), including 60 up-regulated genes and 19 down-regulated genes by the deletion of *recA2* (**Fig. 9A**; details refer to **Table S3**).

Gene ontology (GO) enrichment analysis based on the KEGG database showed that the differentially expressed genes (DEGs) were assigned to 30 GO terms in the categories for biological process, cellular component and molecular function (**Fig. 9B**). Obviously, the biological process DEGs formed the largest group, including 17 GO terms, followed by molecular function (10 GO terms) and cellular component (3 GO terms) (**Fig. 9B**). The DEGs were mainly enriched in two functional regions. One is related to transport and location, including the categories of transport (14 genes), localization and establishment of localization (28 genes), transmembrane transport (5 genes) and protein transmembrane transport (3 genes). The other is related to antioxidantion, including the categories of oxidoreductase activity (3 genes), peroxiredoxin activity (2genes), ferric iron binding (2 genes), antioxidant activity (3 genes) and catalase (1 gene). These DEGs were significantly enriched in ABC transporters and several metabolisms related pathways, such as methane metabolism, biosynthesis of secondary metabolites, metabolic pathways (**Fig. 9C**), but none was in the DNA replication and repair pathways. Combined with the experimental results present in this study, we proposed that the function of *recA2* was mainly related to the gene expression regulation for cellular transportation and antioxidation, which is required for the normal growth of cell.

## Discussion

RecA is an ATPase recombinase reported to play functions in DNA homologous recombination and activation of the LexA-dependent SOS response. Although the *recA* gene is duplicate in some bacterial cells, their functions have less been investigated. In *M. xanthus* DK1622, the expression of *recA1* is very low, which is less than one-tenth of that of *recA2*. The two *recA* genes are both inducible by UV irradiation; but the *recA2* induction is in the early stage, while *recA1* is induced in the late stage. Generally, the gene products expressed in the early and late stages of SOS are responsible for the repair processes and error-prone DNA synthesis, respectively [36, 37]. Thus, the two RecA proteins are both involved in the UV resistance, probably for different damages caused by UV irradiation; RecA2 involves in the early repair processes, and RecA1 is for serious DNA-damage repair, *i.e.* post-replication repair. The deletion of *recA2* caused the mutant to have a long lag phase, but the *recA1* deletion had no significant effect on cellular growth. It is known that the growth lag phase is an adaptation period of bacterial cells to new environment for the changes of temperature and nutrients [38, 39], repair of macromolecule damage and protein misfolding accumulated during cell arrest [40–43], and enzyme preparation for rapid growth in logarithmic phase [38, 42]. Thus, RecA2, instead of RecA1, plays a crucial role in the repairing process required for cellular growth. Consistent to the classic bacterial RecA,

RecA1 possesses the DNA recombination activity and the SOS-gene induction ability, which are required for the survival under the UV irradiation. RecA2 has lost the HR and SOS-gene induction abilities, but has evolved in the gene expression regulation for cellular growth, as well as the cellular survival under the oxidation pressure by hydrogen peroxide. This is the first time to clearly determine the divergent functions of duplicated *recA* genes in bacterial cells.

Amino acid sequence alignment showed that RecA1 and RecA2 amino acid sequences are highly similar in the core ATPase domain, and mainly varied in the N- and C-terminal domains. Lys23 and Arg33 in the N-terminal region are both necessary for the nucleoprotein filament of RecA-ssDNA to capture the homologous dsDNA [28]. The corresponding amino acids at the two sites are both alkaline arginines in RecA1, which is consistent with that in EcRecA. In RecA2, however, the amino acids at the two sites are arginine and proline, respectively (**Fig. 1A**). We aligned the N-terminal sequences of 11 reported bacterial RecAs. The amino acids at the corresponding 23rd site of EcRecA are all Lysine, except Arginine in RecA1 (both are alkaline amino acids), but are less conservative at the 33rd site (**Fig. S1**). Nine RecAs, including RecA1 of *M. xanthus*, are an alkaline amino acid (Arg or Lys) at the 33rd site, while three RecAs did not have the alkaline amino acid there, including RecA2 from *M. xanthus*, RecA from *Prevotella ruminicola* [44] and RecA from *Borrelia burgdorfer* [26]. RecAs with an alkaline amino acid at the 33rd site all have the DNA recombination activity [45–52]. However, similar to RecA2, RecAs from *P. ruminicolah* and *B. burgdorfer* were reported to have no anti-ultraviolet radiation ability [26, 44]. Thus, the lack of an alkaline amino acid at the 33rd site inactivates the DNA recombination activity of RecA enzymes. However, the results present in this study demonstrated that RecA2 of *M. xanthus* is evolved to regulate the genes for cellular transportation and antioxidation, which is obviously related to the damage repair for cellular growth.

As in *E. coli* RecA [35, 53], RecA1 and RecA2 both have the conserved LexA binding sites in their C-terminal regions, including Gly229 and Arg243, and 10 neighboring amino acids (**Fig. 1A**), which, however, does not explain the difference in promoting LexA autolysis between the two proteins. Notably, while the three domains of EcRecA are all acidic, the N-terminal domain of RecA1 and the ATPase domain of RecA2 are alkaline, with the pI values of 9.82 and 8.40, respectively. Accordingly, RecA1 forms more negative charges on the outer side of the polymer, while RecA2 forms more negative charges in the inner side of the helical structure (**Fig. S2**). In adition, unlike the *E. coli* LexA (EcLexA) protein, which is an acidic protein (theoretical pI is 6.23), *M. xanthus* LexA (MxLexA) is a basic protein and its theoretical pI is 8.77. EcLexA and MxLexA are highly conservative, and the difference between the two proteins lies mainly in the linker region (**Fig. S3**). EcLexA linker contains more acidic amino acids (Theoretical pI = 3.58), while the linker of MxLexA contains more basic amino acids (Theoretical pI = 8.75). Besides, MxLexA has two more fragments flanking the linker sequence. The additional fragment at the N-terminal side destroys the β2 folding structure and further lengthens the irregular linker of MxLexA, leading to long irregular chain containing more basic amino acids (Theoretical pI=12.01). According to the binding mode between EcLexA and EcRecA [35], the linker region of LexA is close to the inner groove of the RecA protein filament (**Fig. S4**). The inner helix part of RecA2 (ATPase domain) is alkaline (**Fig. S2B**), which hinders the MxLexA binding to RecA filament and thus hinders its promotion on MxLexA self-cleavage.

Myxobacteria has a relatively large genome size (9-14 Mbp) and contains many DNA repeats [54–56]. These repetitive DNA fragments are potential substrates for RecA-catalyzed homologous recombination. Functional divergence of duplicate RecAs and low expression of the recombination enzyme RecA1 reduce the DNA recombinant activity without affecting other cellular repair functions in *M. xanthus* (RecA2 has no recombination ability but in relatively high expression). In the sequenced myxobacterial genomes (**Table S4**), all the strains, except *Anaeromyxobacter*, have big-size genomes and harbor two *recA* genes. The *Anaeromyxobacter* strains have a single *recA* gene in their genomes, which, however, are in small size (5.0-5.2 Mbp) and possess few repetitive sequences. We propose that functional divergence and expression regulation of duplicate RecAs might be a strategy for the myxobacteria with a large number of repetitive sequences in their big genomes to avoid incorrect recombination.

## Materials and methods

### Strains, media and DNA substrates

Bacterial strains and plasmids used in this study are described in **Table S1**. The *E. coli* strains were routinely grown on Luria-Bertani (LB) agar or in LB liquid broth at 37 °C. The *M. xanthus* strains were grown in CYE liquid medium with shaking at 200 rpm, or grown on agar plates with 1.5% agar at 30 °C [20]. When required, a final concentration of 40 µg/ml of kanamycin (Kan) or 100 µg/ml of ampencillin (Amp) was added to the solid or liquid medium.

The single-stranded viral DNA was isolated from M13mp18 and its 3kb linear dsDNA was amplified by PCR and purified by DNA purification kit (Tiangen). A 60-nt oligomer from M13 genome, 5′-CTG TCA ATG CTG GCG GCG GCT CTG GTG GTG GTT CTG GTG GCG GCT CTG AGG GTG GTG GCT-3′ was synthesize from Tsingke Biotech (Qingdao). The 60-nt oligomer was ^γ-32^P-labeled using polynucleotide kinase [21] and stored in TE buffer (10 mM Tris-HCl, pH 7.0, and 0.5 mM EDTA).

### Growth and resistance analysis

*M. xanthus* strains were grown in CYE medium with shaking at 200 rpm and 30°C to optical density at 600 nm (OD_600_) of 0.5. Cells were then collected by centrifugation at 8000 rpm for 10 min, washed with 10 mM phosphate buffer (pH7.0), and then diluted to 1 OD_600_ in the same buffer.

For radiation damage assay, cells in 10 mM phosphate buffer (pH 7.0) were irradiated at room temperature with a gradient dose from 0 to 200 J/m^2^ using a UV Crosslinker (Fisher Scientific). Then, the cells were re-suspended in fresh CYE medium and incubated at 30 °C for 4h. After post-incubation, cells were harvested by centrifugation, and used for further assay or stored at -80 °C.

For oxidative damage assay, cells were suspended in a phosphate buffer (pH 7.0) with a concentration of 1 OD, and hydrogen peroxide (H_2_O_2_) was added to the final concentration from 1 to 5 mM. The bacterial suspension was incubated for 20 min at room temperature with gentle shaking. After treatment, the suspension was immediately 10-fold diluted in the same phosphate buffer to end oxidative damage reaction. Then, cells in the suspension were collected for further assay.

The growth assay was determined by growing in a liquid medium at 30 °C. Strains were inoculated at 0.02 OD_600_ and grown in CYE media for 84 h with shaking at 200 rpm. OD_600_ was read every 12 h.

The survival assay was determined by soft agar colony formation assay. Briefly, 15 ml CYE solid medium in 9-cm culture dish was used as bottom layer. Strains were diluted with fresh medium, mixed at the 1:2 ratio with melted 0.6% soft agar (50 °C), and put the mixture into the CYE plates. After a few minutes for medium solidification, the cultures were incubated until clone formation. The survival percentage was calculated as the number of colony-forming unit (CFU) (damaged) divided by the total number of CFU (Undamaged).

### Genetic manipulations

*E. coli* Plasmids were isolated by the alkaline lysis method and the chromosomal DNA of *E. coli* or *M. xanthus* was extracted using bacterial genome DNA extraction kit (Tiangen). Cloning of genes including *recA1*, *recA2*, and *lexA* from *M. xanthus* were operated according the general steps [21]. The genes were amplified by PCR and was ligated into the expression plasmid pET15b, respectively. The primers used here are listed in **Table S2**.

Mutant construction was performed using the markerless mutation in *M. xanthus* DK1622, with the pBJ113 plasmid using the kanamycin resistant cassette for the first round of screening and the *galK* gene for the negative screening [22]. Briefly, the up- and down-stream homologous arms were amplified with primers (listed in **Table S2**) and ligated at the *Bam*H1 site. The ligated fragment was inserted into the *Eco*RI/*Hin*dIII site of pBJ113. The resulting plasmid was introduced into *M. xanthus* via electroporation (1.25 kV, 300 W, 50 mF, 1 mm cuvette gap). The second round of screening was performed on CYE plates containing 1% galactose (Sigma). The *recA1* mutant (named RA1) and *recA2* mutant (named RA2) were identified and verified by PCR amplification and sequencing.

### RNA extraction, RT-PCR and RNA-Seq assay

Total RNA of *M. xanthus* cells was extracted using RNAiso Plus reagent following the manufacturer’s protocol (Takara, Beijing). The cDNA synthesis used the PrimeScript RT Reagent Kit with random primers. The synthesized cDNA samples were diluted 5 times prior to RT-PCR. The primers were designed for *lexA*, *recA*1 and *recA*2 (**Table S2**). RT-PCR was accomplished using the SYBR premix Ex Taq kit (Takara, China) on an ABI StepOnePlus Real-Time PCR System (Thermofisher Scientific, USA). Gene expression was normalized with the *gapA* expression and calculated using the equation: change (x-fold) = 2^−ΔΔCt^ [23].

RNA sequencing was conducted in Vazyme (Beijing). Three independent repeats are set for each sample. All the up-regulated and down-regulated genes were obtained by comparing with control, and their gene functions were annotated using the NR, GO, and KEGG databases.

### Protein expression, purification and characterization

The constructed expression plasmid with *recA1*, *recA2*, or *lexA* was introduced into *E. coli* BL21(DE3) competent cells. The protein expression was induced with 1mM IPTG and purified with Ni-NTA agarose according the manual of Ni-NTA purification system (Invitrogen). After overnight dialysis with storage buffer (20mM Tris, 150mM NaCl, 0.1mM DTT, 0.1mM EDTA, 50% glycerol), the purified proteins were quantified and stored at -80 °C.

ATPase activity of RecA protein was determined in the presence or absence of DNA according to the methods described previously [24]. Final reaction mixture in a 2-ml volume contained: 20 mM Tris-HCl (pH 7.4), 10 mM NaCl, 5 mM MgCl_2,_ 2 mM KCl, 3 mM ATP (Sigma), 1 mM CaCl_2_, 1 mM DTT, and 2% glycerol. The mixture was preheated to 32 °C before the addition of RecA and DNA. ATPase activity was determined by measuring the free phosphate ion (Pi) released from ATP using the ultramicro ATPase activity detection kit (Nanjing Jiancheng Bioengineering, Nanjing).

*In vivo* LexA cleavage analysis was performed as described previously [24].

D-loop assays for strand assimilation were performed according to the previously described methods [25, 26] with some modifications. Briefly, 0.2 µM RecA and 10 nM ^32^P-labeled ssDNA was combined in 9 µl of reaction buffer containing 25 mM Tris-HCl (pH 7.5), 75 mM NaCl, 5 mM MgCl_2_, 3 mM ATP, 1 mM DTT, 1 mM CaCl_2_, and incubated at 37 °C for 5 min. Then 1 µl of RF M13 plasmid was added to the final concentration of 1 μM and incubation at 37 °C was continued for 20 min. The reaction was stopped by adding sodium dodecyl sulfate to 0.5% and proteinase K to 1 mg/ml. The deproteinated reaction products were run on a 0.9% agarose 1× TAE gel and visualized using autoradiography with phosphor screen.

## Acknowledgements

This work was financially supported by the National Natural Science Foundation of China (NSFC) (Nos. 31670076 and 31471183), the National Key Research and Development Program (2018YFA0901700), and the Key Program of Shandong Natural Science Foundation (No. ZR2016QZ002) to YZL.

The funders had no role in study design, data collection and analysis, decision to publish, or preparation of the manuscript.

## Competing interests

The authors declare that they have no competing interests.

---

**Figure S1.**
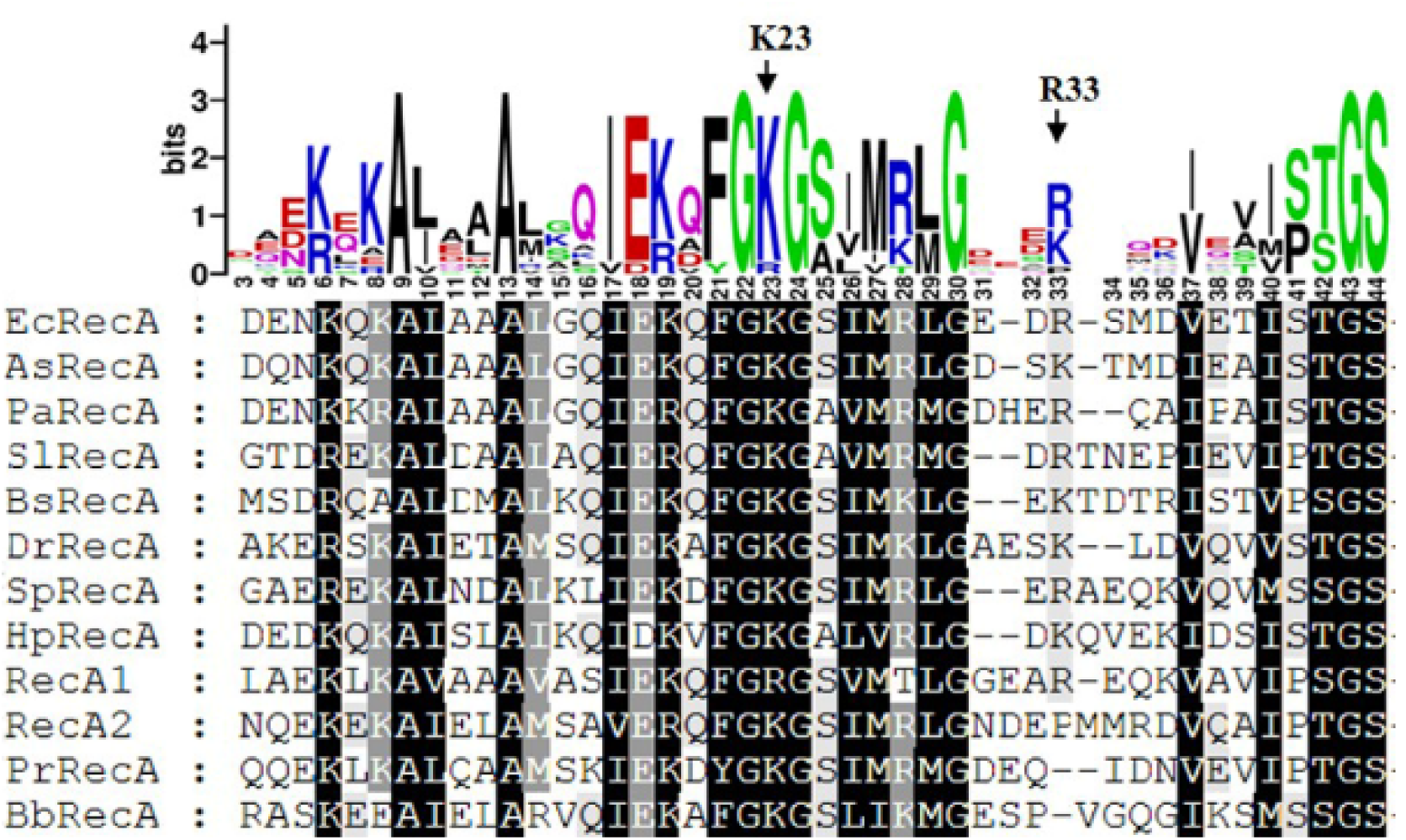
Amino acid sequence alignment of RecA N-terminal domains from different bacteria, including EcRecA from *E. coli* (b2699) (Cox, 1999); AsRecA from *Aeromonas salmonicida* (ASA_3809) (Umelo et al., 1996); PaRecA from *Pseudomonas aeruginosa* (PA3617) (Sano et al., 1987); SlRecA from *Streptomyces lividans* (SLIV_09770) (Nussbaumer et al., 1994); BsRecA from *Bacillus subtilis* (BSU16940) (Carrasco et al., 2019); DrRecA from *Deinococcus radiodurans* (DR_2340) (Kim et al., 2002); SpRecA from *Streptococcus pneumoniae* (SP_1940) (Grove et al., 2012); HpRecA from *Helicobacter pylori* (HP0153) (Orillard et al., 2011); PrRecA from *Prevotella ruminicola* (PRU_0066) (Aminov et al., 1996); BbRecA from *Borrelia burgdorferi* (BB_0131) (Huang et al., 2017). The key amino acids are noted (black arrow), and numbering is shown based on that of *E. coli* RecA.

**Figure S2.**
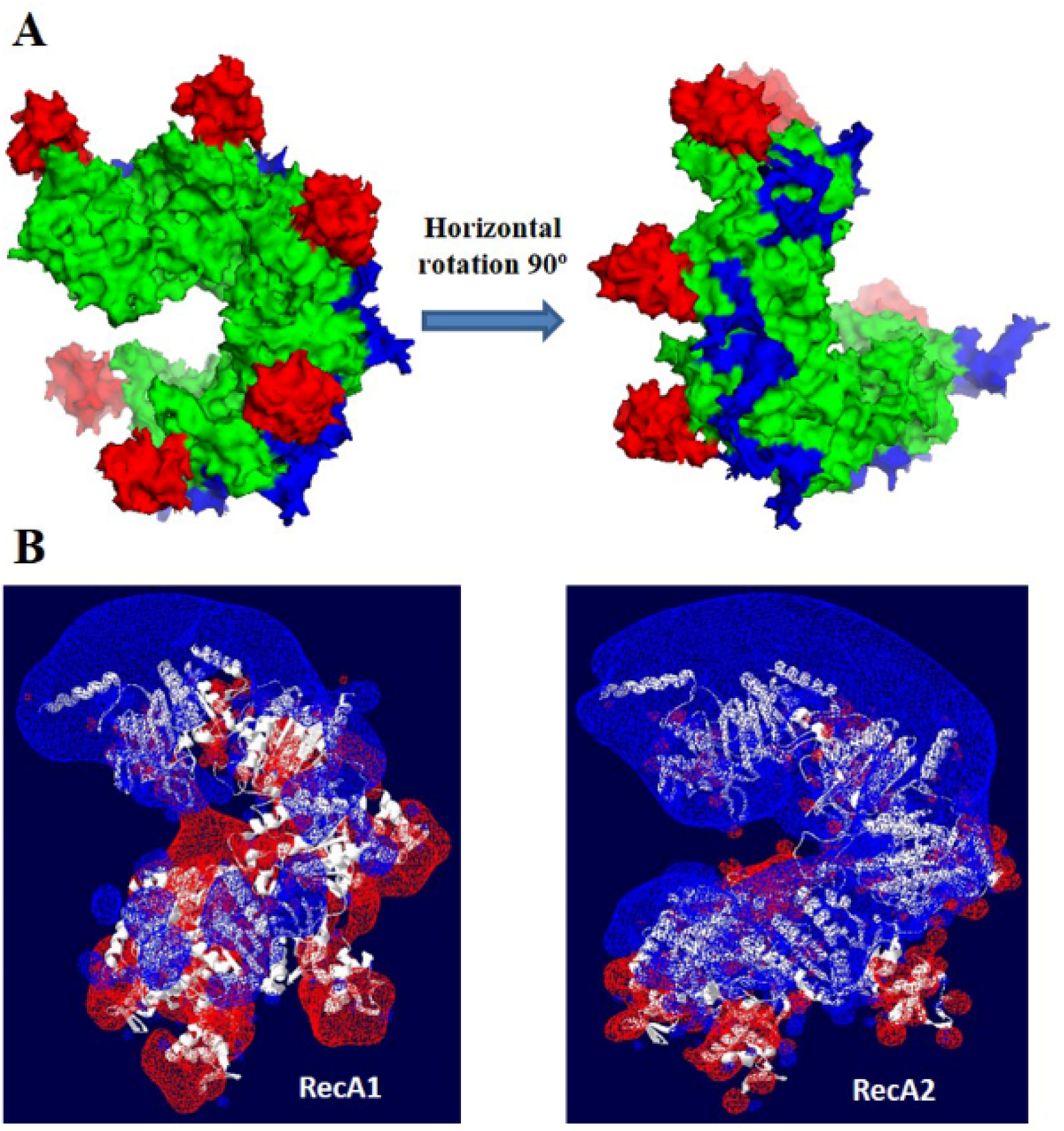
Charge analysis on RecA1 and RecA2. (A) Locations of NTD (red), CTD (blue) and the core ATPase domain (green) in the RecA polymer. In the helical polymer model of RecA proteins, the N- and C-terminal structures are both in the outer side, and the ATPase domain is in the inner side of the polymer. The right one is the left one flipping 90 degrees horizontally. (B) Surface charge of RecA1 and RecA2. Blue represents negative charge and red represents positive charge.

**Figure S3.**
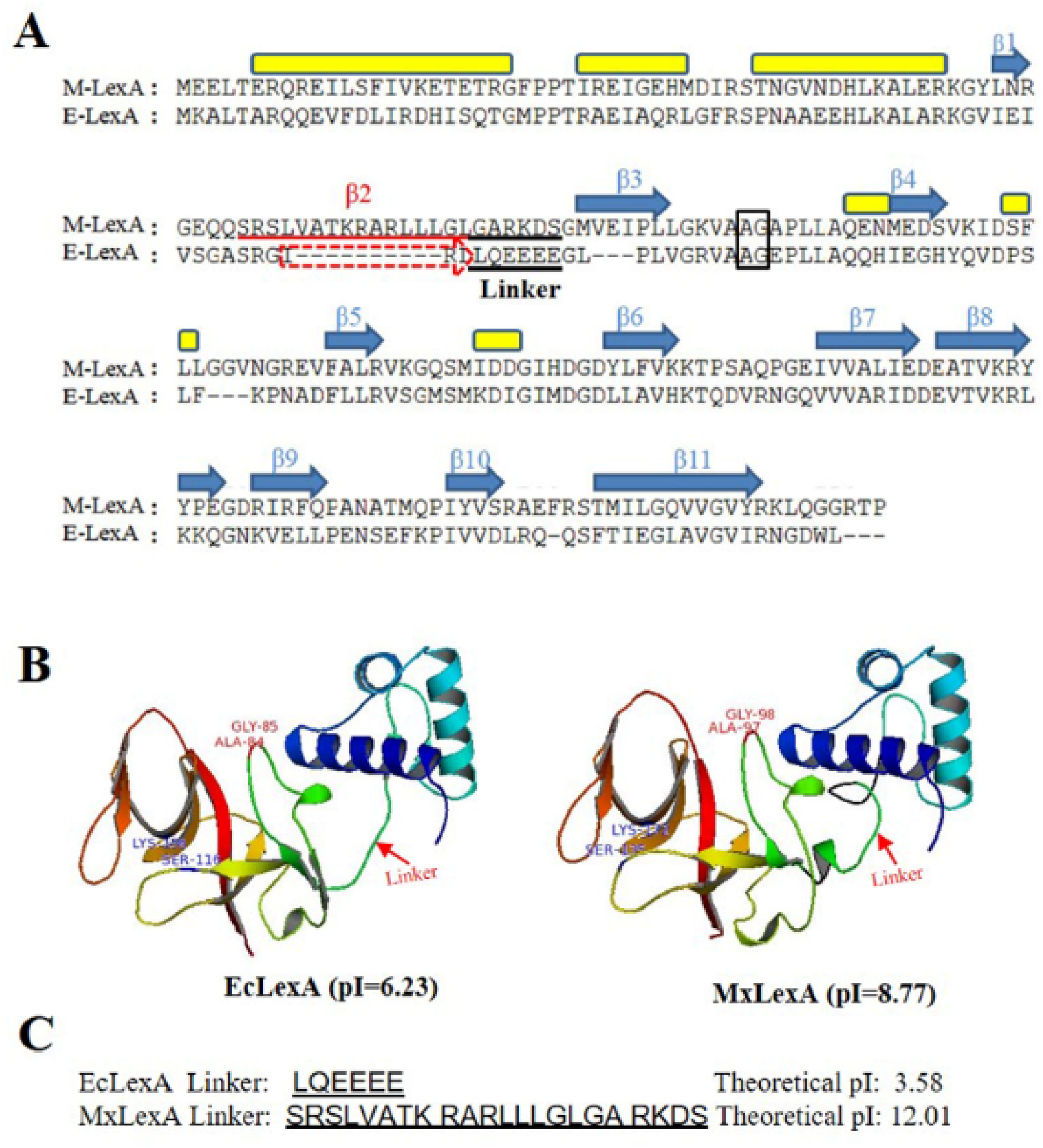
Sequence and structural comparison of the LexA proteins from *M. xanthus* and *E. coli*. (A) Sequence alignment of *M. xanthus* LexA (MxLexA) and *E. coli* LexA (EcLexA) using the MUSCLE program. The LexA self-cleavage site A-G was marked in a black box. The linkers between the N- and C-terminals of the two LexAs are underlined. (B) The 3D structure of MxLexA was constructed using homology modeling with EcLexA PDB structure (1JHF) as template. The linkers between the N- and C-terminals of the two LexAs are indicted with red arrow. (C) The sequences and theoretical pI values of the linkers of EcLexA and MxLexA.

**Figure S4.**
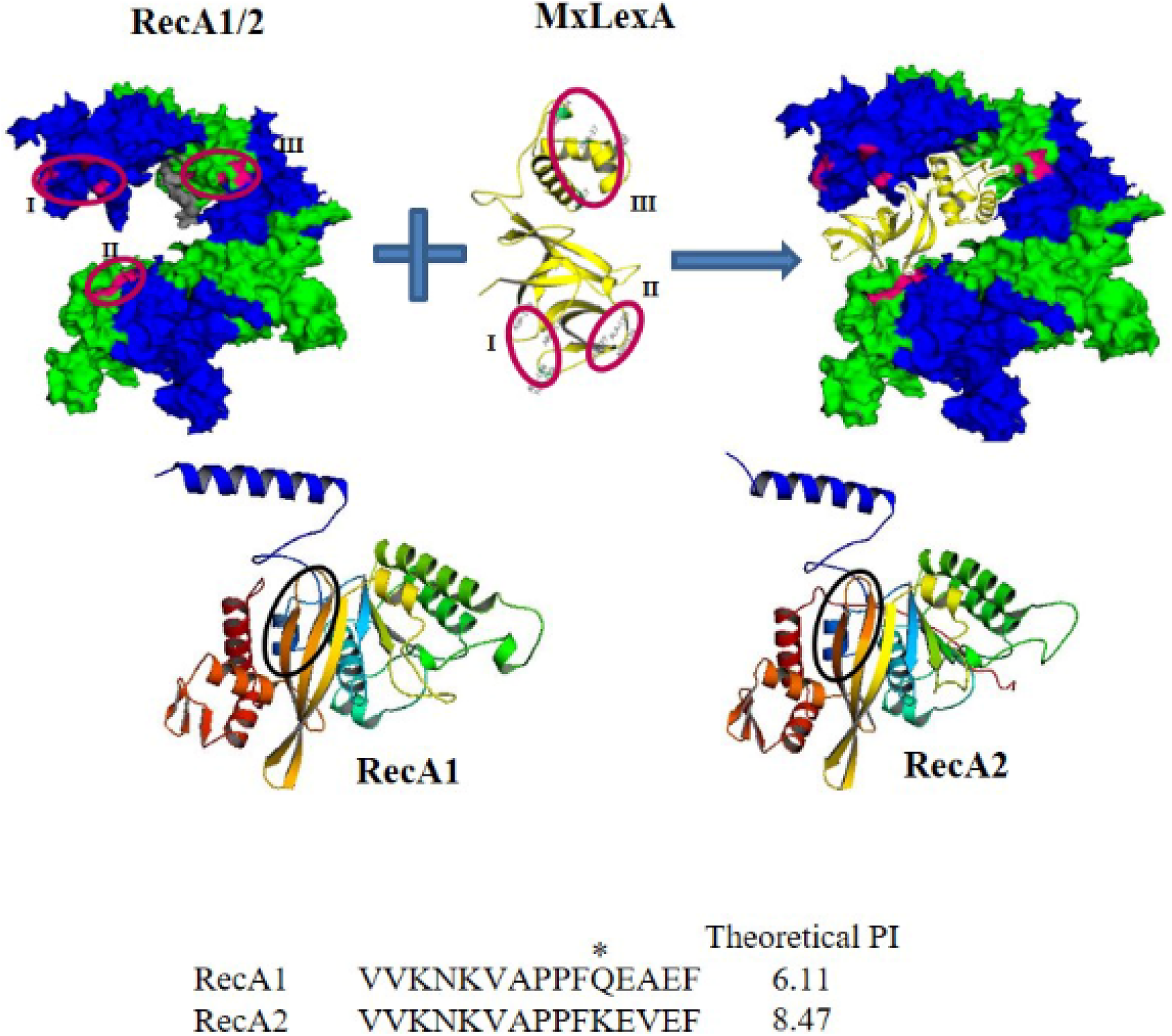
Simulated docking of LexA and RecA polymers. LexA and RecA bind to each other mainly through three binding regions (Kovačič et al., 2013), which are marked in red circle (upper map). The possible binding region of the Linker of LexA on RecA polymer is marked with grey and marked with black circles in RecA1 and RecA2 (lower map). The amino acid sequence of the linker binding regions and their corresponding PIs are listed below the figure.

**Figure S5.**
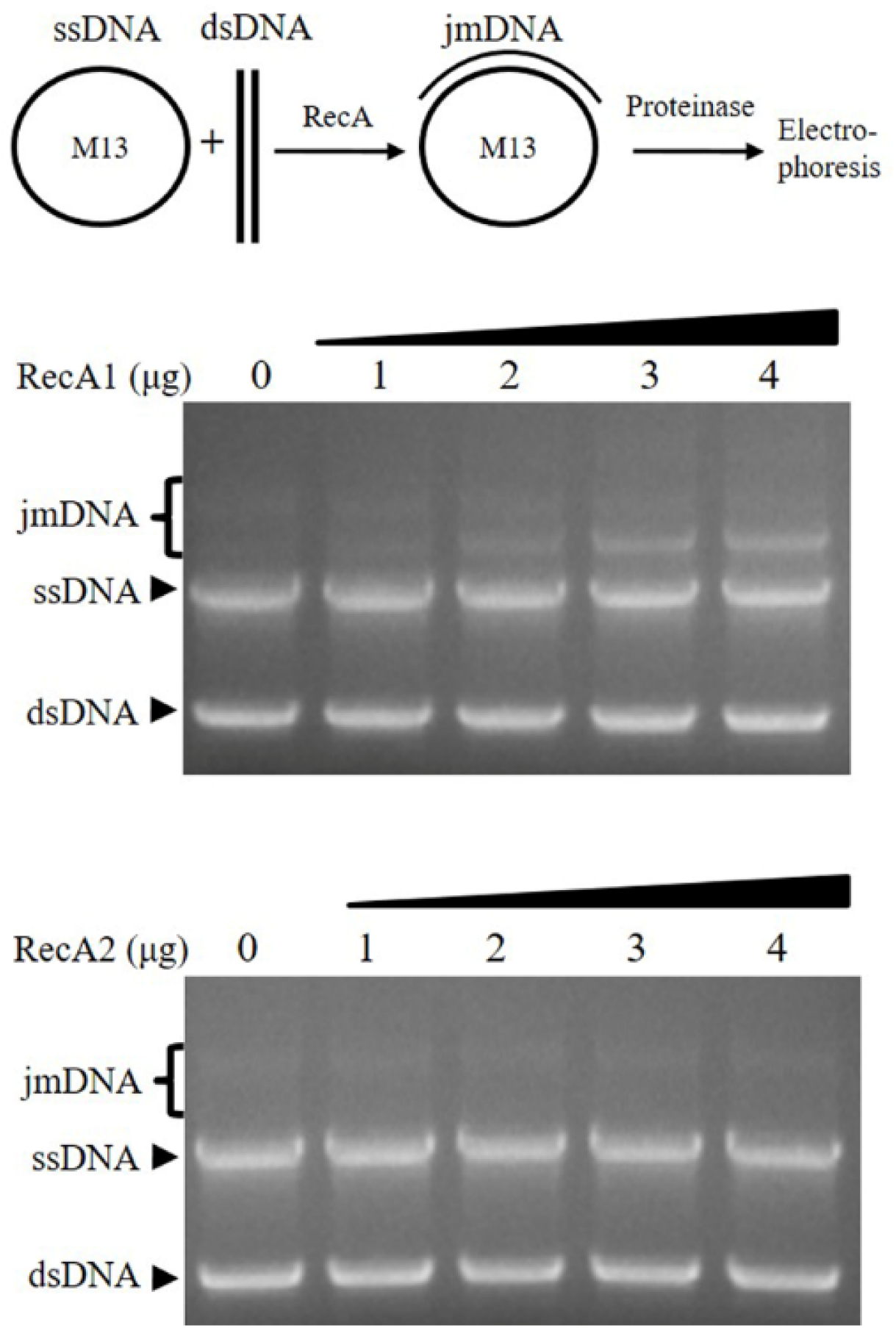
DNA strand exchange reaction promoted by RecA between M13 circular ssDNA and the linear dsDNA (derived from M13). Reactions were performed at 30 °C in a solution containing 25 mM Tris-HCl, pH 7.0, 50 mM NaCl, 4% glycerol, 1 mM DTT, 10 mM MgCl_2_, 3 mM ATP and an ATP-regenerating system (10 units/ml of pyruvate kinase/3.3 mM phosphoenolpyruvate). After pre-incubation of ssDNA with RecA1 or RecA2 protein at 30 °C for 5 min, linear duplex DNA was added to start the DNA strand exchange reactions. The ssDNA and dsDNA substrates, as well as the joint molecule intermediates (jmDNA) bands are all visible in the 0.8% agarose gel.

**Table S1.** Strains and plasmids used in this study

| Strains or plasmids | Genotype or description | Source or references |
| --- | --- | --- |
| **Strains** |  |  |
| *M. xanthus* |  |  |
| **DK1622** | Wild-type strains | D. Kaiser, University of Stanford |
| **RA1** | DK1622 (ΔrecA1) | This study |
| **RA2** | DK1622 (ΔrecA2) | This study |
| *E. coli* |  |  |
| **DH5α** | F<sup>-</sup>80dlacZΔM15 $\Delta$(lacZYA<sup>-</sup>argF)U169 deoR recA1 endA1 hsdR17 (r<sub>k</sub><sup>-</sup>m<sub>k</sub><sup>+</sup>) phoA supE44 $\lambda$ thi-1 gyrA96 relA1 | Takara |
| **JM109** | recA1, endA1, gyrA96, thi-1, hsdR17, supE44, relA1, $\Delta$(lac<sup>-</sup>proAB)/F'[traD36, proAB<sup>+</sup>, lacIq, lacZΔM15] | Promega |
| **BL21(DE3)** | F<sup>-</sup>lon ompT hsdSB (rB<sup>-</sup>, mB<sup>-</sup>) dcm gal dcm(DE3) | This study |
| **BL2/pET15recA1** | Expression strain of recA1 with plasmid pET15recA1 | This study |
| **BL21/pET15recA2** | Expression strain of recA2 with plasmid pET15recA2 | This study |
| **Plasmids** |  |  |
| **pBJ113** | Gene replacement vector with KG cassette, Kan<sup>r</sup> | Z.M. Yang, Virginia Tech |
| **pBJ-recA1** | Upstream and downstream homologous arms of recA1 inserted into EcoRI/HindIII site of pBJ113, Kan<sup>r</sup> | This study |
| **pBJ-recA2** | Upstream and downstream homologous arms of recA2 inserted into EcoRI/HindIII site of pBJ113, Kan<sup>r</sup> | Wu & Kaiser, 1995 |
| **pET15recA1** | Recombination plasmid with a recA1 gene inserted into NdeI/BamHI sites of pET15b, Amp<sup>r</sup> | This study |
| **pET15recA2** | Recombination plasmid with a recA1 gene inserted into NdeI/BamHI sites of pET15b, Amp<sup>r</sup> | This study |
| **pET15lexA** | Recombination plasmid with a lexA gene inserted into NdeI/BamHI sites of pET15b, Amp<sup>r</sup> | This study |
| **M13mp** | a filamentous E. coli bacteriophage, used for DNA recombination assay | NEB |

**Table S2.**
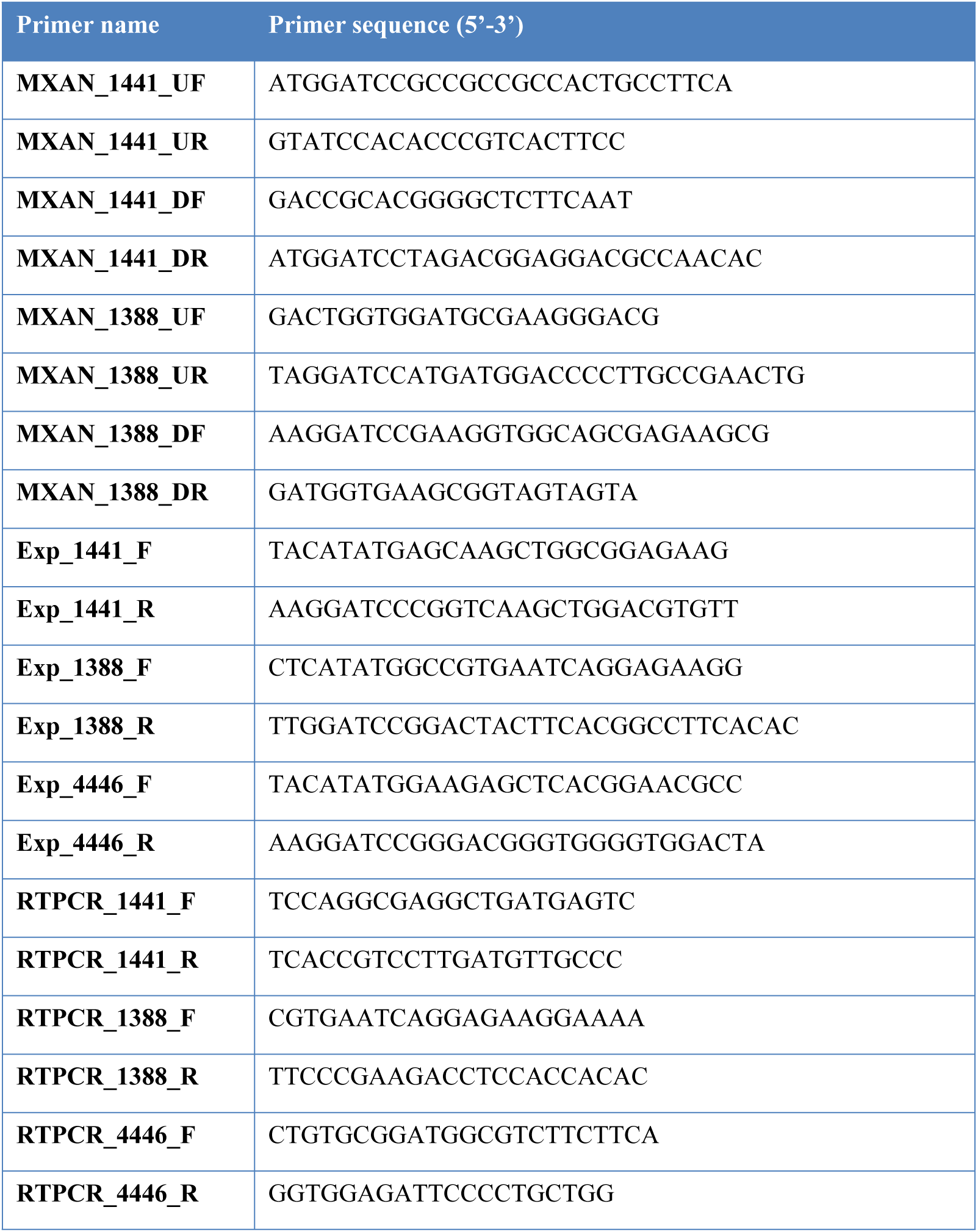
Primers used in this study

**Table S3.**
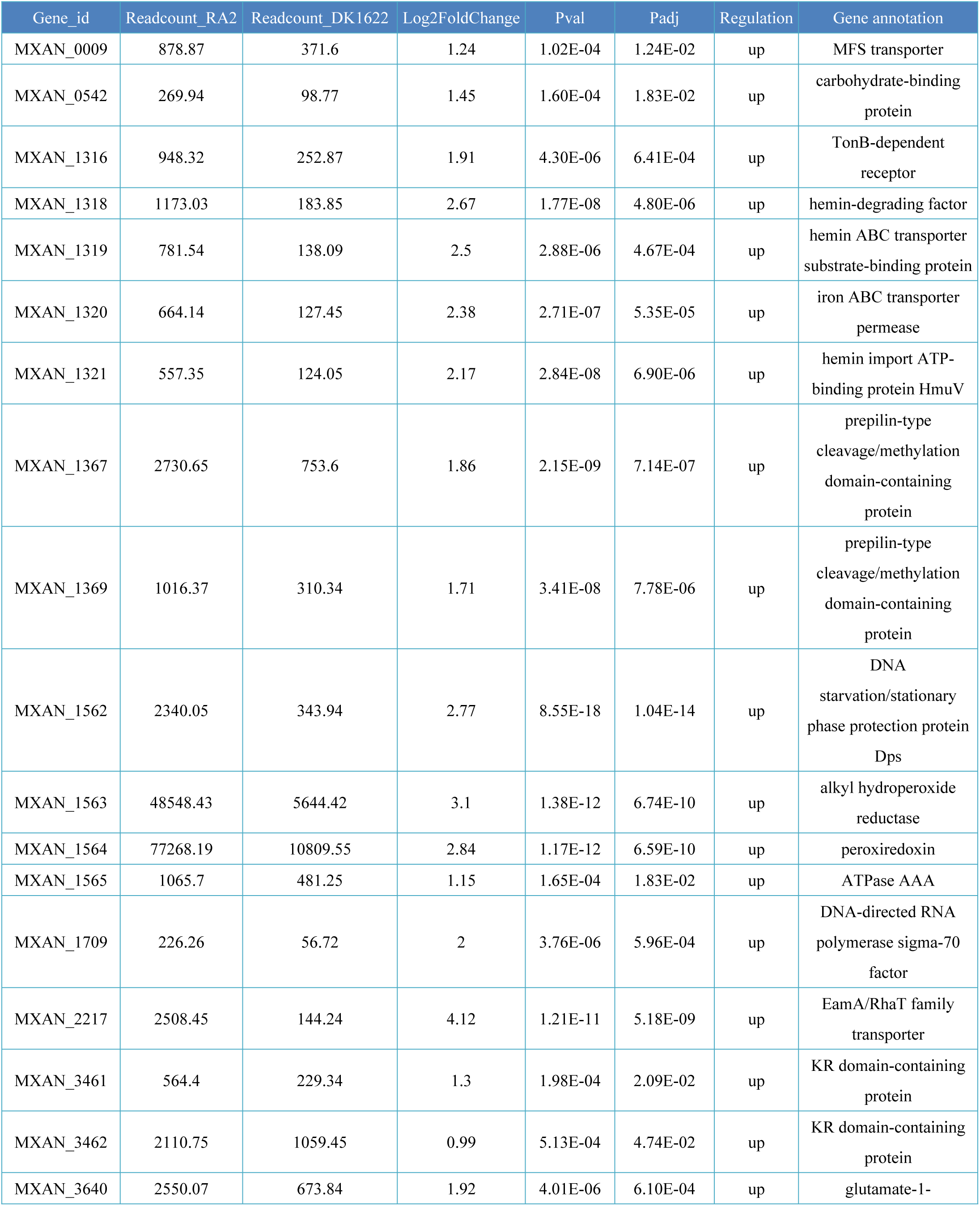
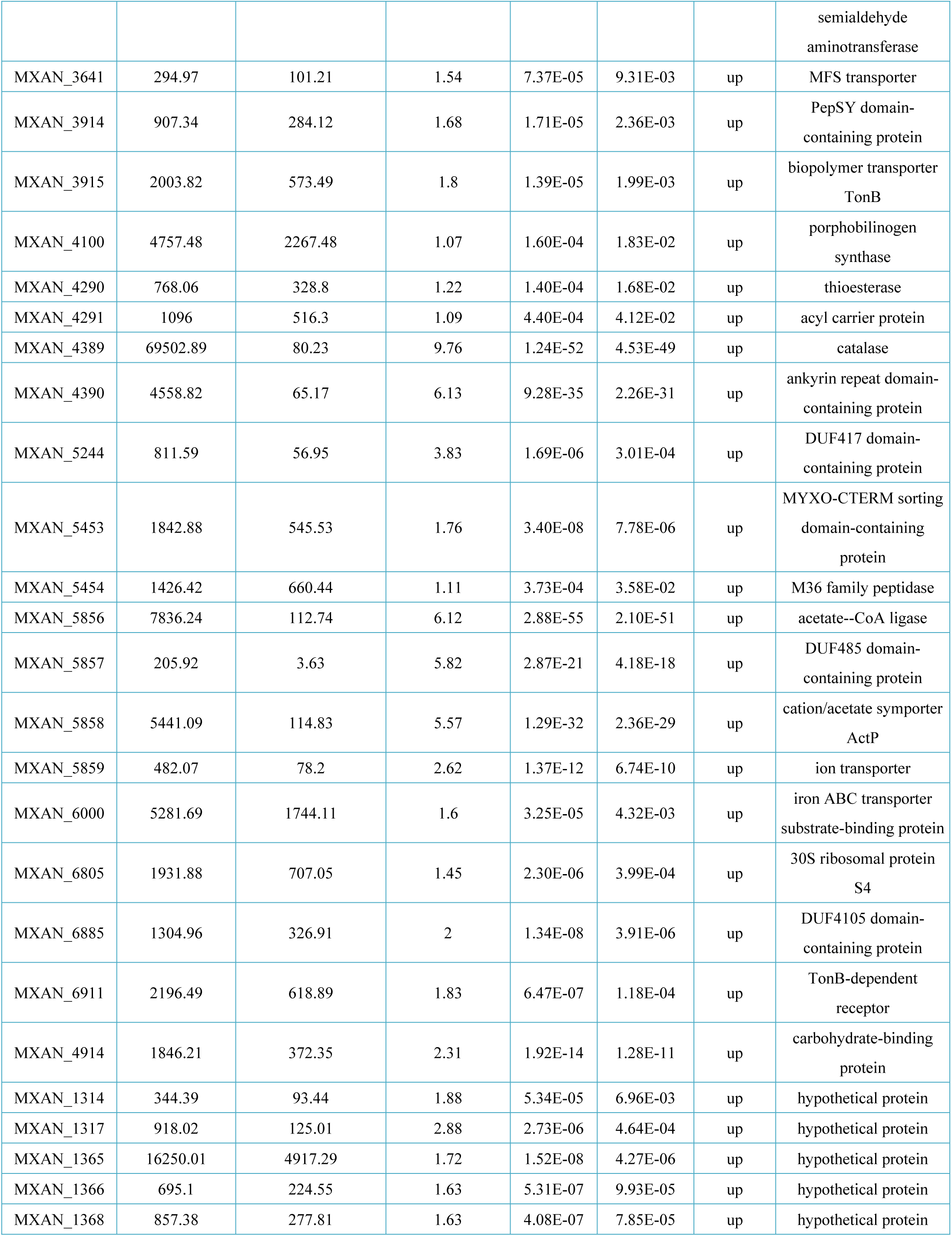
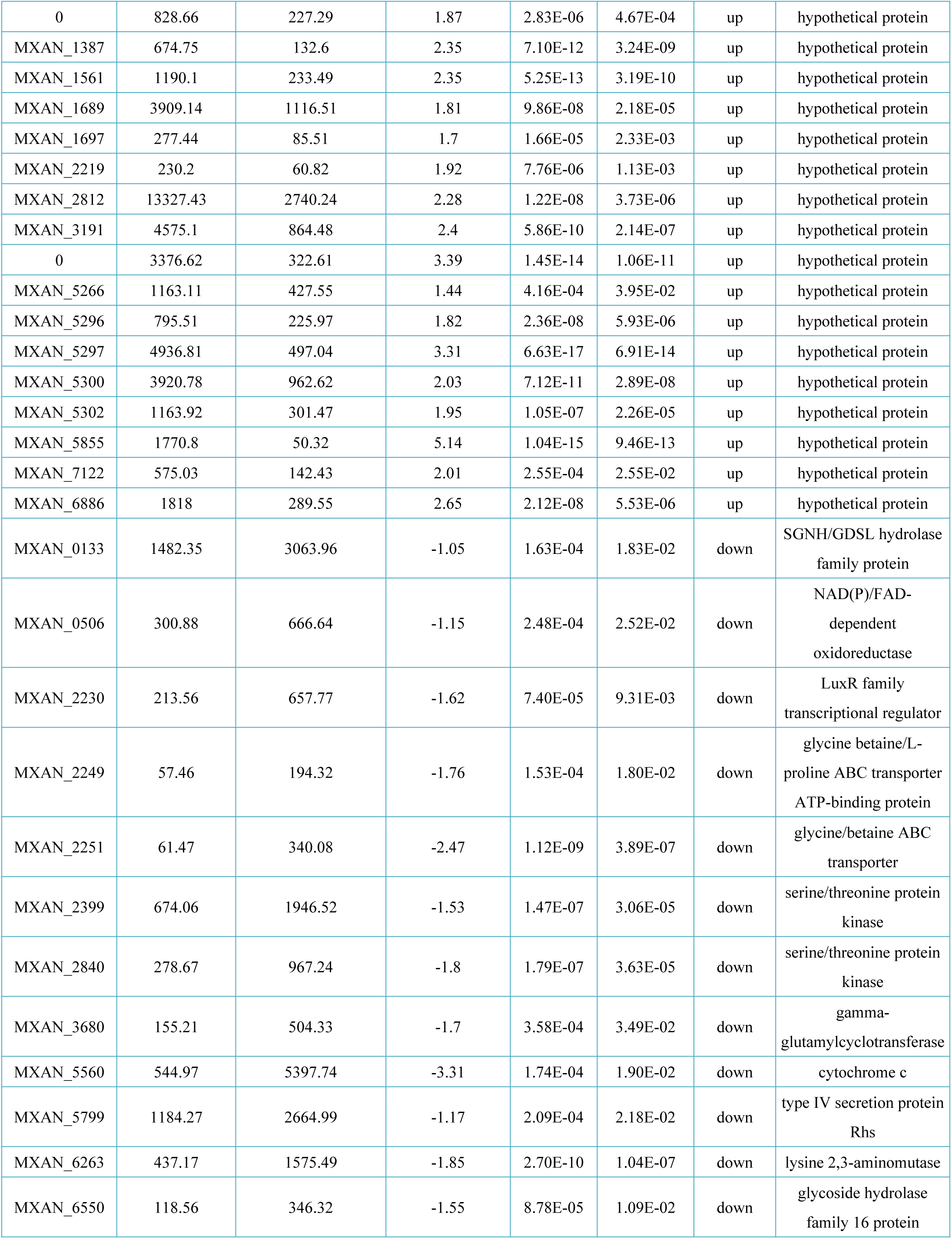
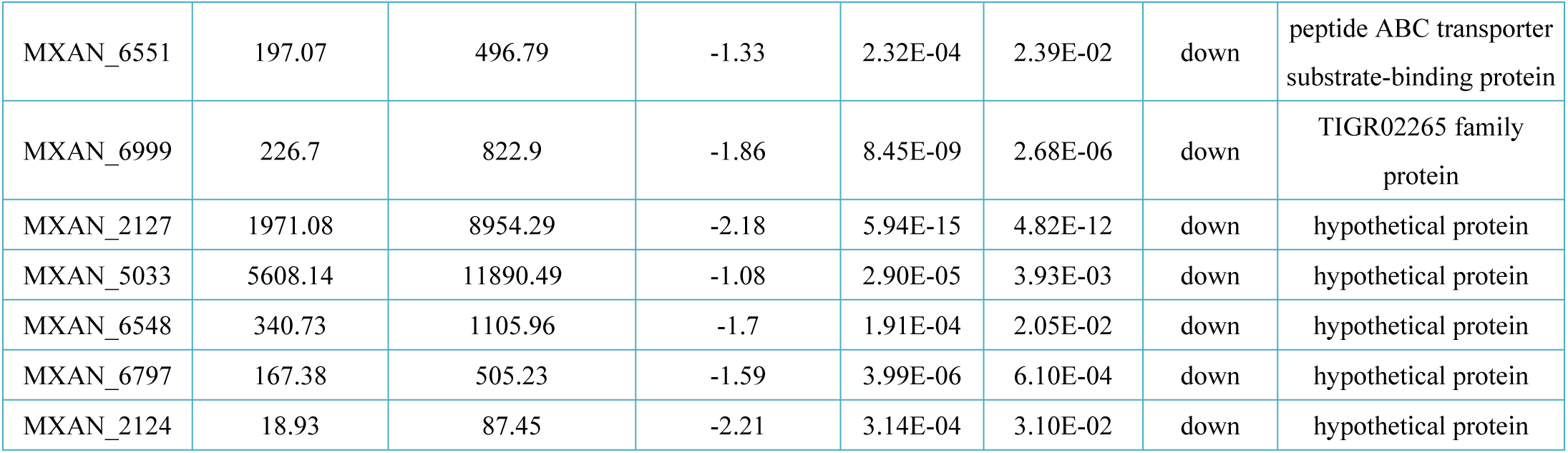
List of differentially expressed genes between the transcriptomes of RA2 and DK1622

**Table S4.**
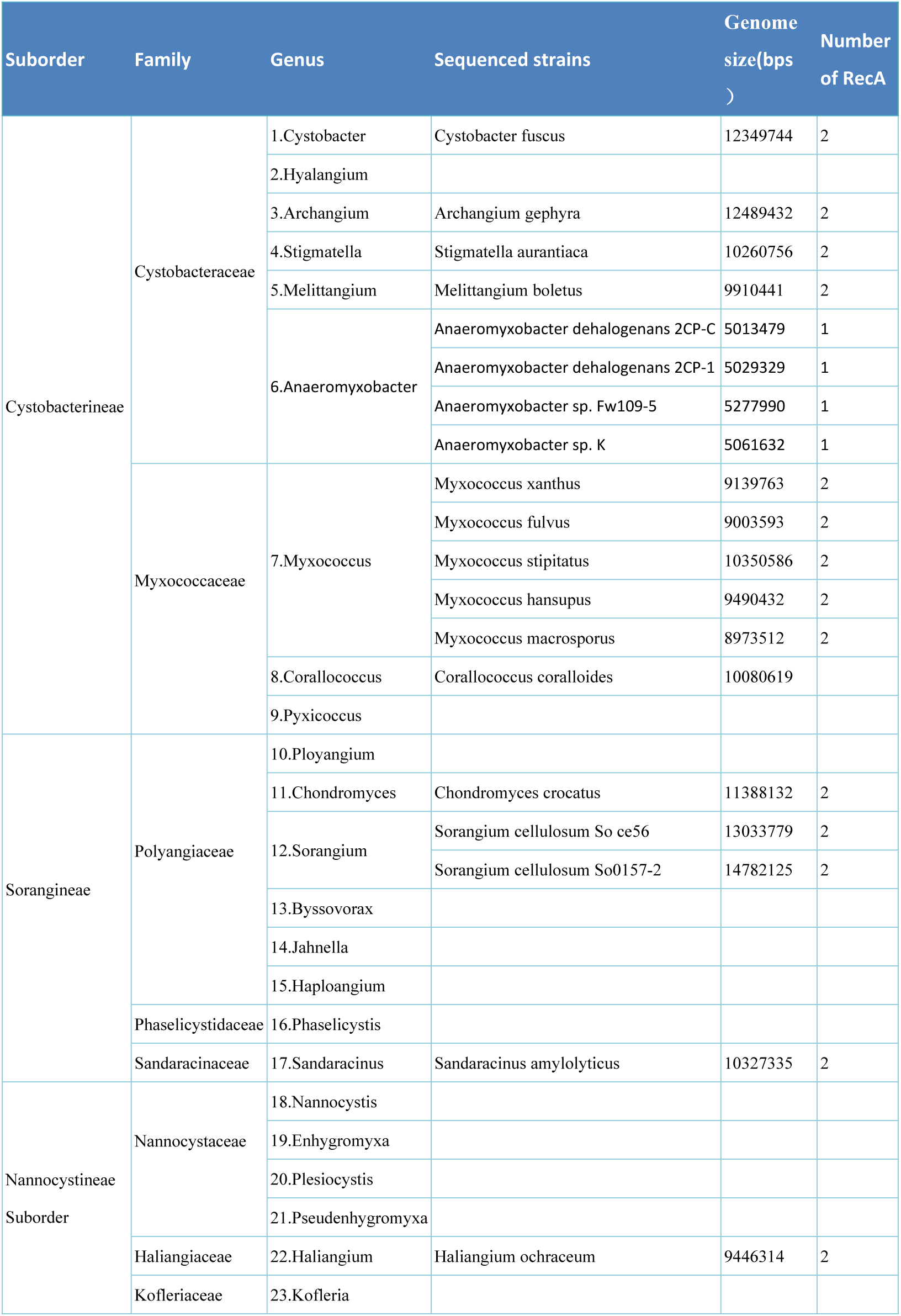
Sequenced myxobacteria genome size and RecA duplication.

